# K^+^ in a K_v_1.2 Channel Pore: Hydration, Selectivity, and the Role of a Conserved Threonine

**DOI:** 10.1101/638783

**Authors:** A. M. Kariev, M. E Green

## Abstract

Quantum calculations describing transport of K^+^ through a K_v_1.2 channel cavity, plus the lower half of the selectivity filter (SF), show hydration in the pore and cosolvation by threonine at the entrance to the SF. Comparison to calculations on Na^+^ ions gives the probable selectivity mechanism. A single K^+^ ion is calculated at five positions in its course through the cavity, and two ions calculated at three positions at the entrance to the SF. Three Na^+^ pairs of ions were also calculated, and one shows how an ion is trapped asymmetrically, tightly held by two threonine −OH, and with a water tightly bound ahead of it, so that overall it has a major barrier to advancing, while K^+^ advances with minimal barriers. In the cavity below the SF, the ion passes in a hydrated state through pore water, between the intracellular gate and the SF, until it is cosolvated by the threonines at the selectivity filter entrance. These calculations show how the ion associates with the water, and enters the SF. A characteristic arrangement of four water molecules adjacent to the SF in the KcsA channel, shown in earlier work, is now found in K_v_1.2. A single ion passing through the channel cavity is found to have an energy minimum within 1 Å of the K^+^ ion position in the 3Lut pdb structure of this channel. Properties (e.g. dipole moment) of the system are calculated. Charge transfer to the ion produces K^+^ charge 0.74 ≤ q(ion) ≤ 0.87 *e*, in different conditions. The calculations of pairs of Na^+^ and K^+^ ions at the SF entrance include the threonine, valine, and glycine of the conserved SF TVGYG sequence. The Na^+^/K^+^ difference shows a reason for the conservation of the threonine in producing selectivity, as the –OH groups trap Na^+^ but not K^+^.

**STATEMENT OF SIGNIFICANCE:** Potassium channels are found in all cells, and have a characteristic selectivity filter that blocks the passage of Na^+^ while allowing K^+^ to pass. These channels are implicated in many diseases. We use quantum calculations to show how the K^+^ ion passes from the intracellular gate of the channel, entering the channel pore, to the selectivity filter at the extracellular end of the channel; at the selectivity filter, we use comparable calculations of K^+^ and Na^+^ to show how the channel selects K^+^ over Na^+^, as well as the probable reason for the conservation of a key residue (threonine) at the base of the selectivity filter. We find properties (e.g., charge transfer, bond order) that require quantum calculations.

## INTRODUCTION

Potassium channels are essentially ubiquitous in biological cells; they are even found in viruses(1, 2). Malfunctions of these channels are responsible for multiple diseases(3–6). To understand their functions, there are four major classes of questions that must be addressed: 1) the mechanism of gating, or opening and closing the channel 2) the mechanism of conduction of ions in the open channel, 3) how the channel stops conducting without returning to the original closed configuration (C-type inactivation) 4) the mechanism of selectivity: why does a K^+^ channel conduct K^+^ >10^3^ times better than it conducts Na^+^?(7) Here we consider questions 2 and 4, conductivity and selectivity. Water is critical in the passage of ions through the pore, as well as in transferring the ion from the pore to the selectivity filter (SF), a relatively narrow region near the extracellular end, generally considered responsible for the selectivity of the channel. There is a cavity, or vestibule, between the activation gate and the SF, and the passage of a potassium ion through this section has not been the focus of as much attention as has been devoted to the other questions. However, this is a necessary part of the conduction path, and the rearrangement of water in this path is a key to the connection of the gate to the SF.

The passage of the ion through the vestibule, or cavity, between the gate and the SF has been studied in a number of ways, including considering the channel as though it were a macroscopic system (8). However, it is necessary to understand the mechanism at atomic level resolution. Conduction has generally, for many years, been described in terms of some variation of a knock-on mechanism, which couples entry of an ion into the gate and on into the cavity, to the progress of an ion already in the cavity on into the SF. Much of the evidence comes from the KcsA channel, a bacterial channel that is gated by a drop in pH, and which does not contain a voltage sensing domain. 2D spectroscopy of the SF of a semi-synthetic KcsA channel favors a knock-on model for ion transport that requires water(9); no model that omits water could be compatible with the data. Kraszewski and coworkers carried out a computation of the positions of the ions in the SF showing a barrier-less transition consistent with a knock-on mechanism, and emphasizing the importance of a charge at the extracellular mouth of the SF. (10) However, KcsA has a relatively short cavity, which appears to have 12 water molecules, while the longer K_v_1.2 has many more.

However, the SF is almost identical in KcsA and the voltage-gated channels. Cuello, Marien Cortes and Perozo(11) determined the KcsA structure in several cases, which they attributed as follows (pdb codes given first): 1k4c (12), closed/open and 1k4d, closed/inactivated (12); 5VK6, open/open and 5VKE, open/inactivated, where the open, closed distinction refers to the activation gate, followed by the open, inactivated designation for the SF(11); inactivation is a state in which the channel ceases to conduct while not returning to the resting state, and is attributed to a kind of constriction of the SF. Some years earlier, we did a quantum calculation, starting from the 1k4c structure, showing the water in the pore under the SF with four molecules hydrogen bonded to the threonines at the bottom of the SF(13), a configuration we called a “basket”. Their experimental structures show the positions of crystallographic water in the pore, and the hydration of the K^+^ ion. Our calculation showed the position of the four “basket” water molecules with the orientation of the hydrogens, and their hydrogen bonding to SF threonines. This holds the water structure together (in addition to the one bond per water with its neighboring water). The calculated water structure shows approximately the correct spacing of the molecules, according to the structural study of Cuello et al on the 1k4c structure, which we used (14, 15). Repeating this for the K_v_1.2 (3Lut structure, with H atoms added (16)) structure, and finding essentially the same “basket”, generalizes this result.

The relation of water to selectivity, particularly the structure at the threonine at the junction of the pore and the SF, has been studied extensively. De et al (17) considered the entire SF, and found that Na^+^ bound more tightly than K^+^ at val76 in KcsA, the residue just above threonine in the SF. However, Krishnan et al (18) showed by mutations that the threonine played the major role in K^+^/Na^+^ selectivity. Noskov et al (19) used MD to argue for protein flexibility in the SF as being important for selectivity. However, Wang et al (20) used FRET, in a somewhat different potassium channel (the inverse rectifier Kir1.1), but with similar SF, to argue that flexibility in some mutants destroys selectivity. Shen and Guo proposed that fluctuations in the SF of a channel mutated from the non-selective NaK channel were important (21). A strong role for electrostatics has also been considered (22). The subject has also been reviewed multiple times (e.g., (23–25)). Given the range of proposals, clearly there is more to be done to elucidate selectivity. However, as far as we are aware no previous quantum calculation has shown the specific structure as the ion moves from the pore into the initial position in the SF. Two previous studies using quantum calculations are particularly relevant, however. Huetz et al (26) carried out a partially QM calculation on the SF. They did not look for asymmetry in the position of Na^+^ but they very carefully studied the charge transfer to the ions. with results very similar to those we find. Their results do not show in detail how Na^+^ and K^+^ differ in their linkages to oxygens in the SF, nor the role of the threonine hydroxyls, but they have the basic calculation that would lead to understanding selectivity. We appear to have carried out the only other previous relevant quantum study, on KcsA (13, 14). We did find the asymmetry of the Na^+^ binding, but did not have the detail that explains selectivity and the importance of the threonine hydroxyls.

Water plays a role in properties other than conduction and selectivity. While we do not consider inactivation in our calculations, it is worth mentioning that inactivation also has a major role for water in and near the SF, as has been discussed by multiple authors. Water is required within the SF for inactivation in the KcsA channel (27). Additional evidence for the importance of water in inactivation is given by Ostmeyer et al (28); Raghuraman et al (29) showed how water becomes more nearly bound, less mobile, at the outer vestibule in the inactivated state. There is evidence for the presence of three water molecules in a crevice behind the SF, which play a key role in inactivation. Li and coworkers carried out MD studies, primarily on the SF(30, 31). Furini et al have used MD to compute the osmotic permeability of a KcsA channel, without ions, in response to a pressure gradient (32). An osmotic pulse method was used by Ando and coworkers (33) to determine the coupling of K^+^ and water motion through the hERG potassium channel. Fowler et al concluded that MD could not give results that allowed comparison with experiments (34); the same group used MD to determine a single transition in the SF (35).

## THIS CALCULATION: *METHODS*

All calculations used Gaussian09(36), versions D or E. Natural Bond Order (NBO)(37) calculations were done under Gaussian also. Optimization was done at HF/6-31G* level, and the optimized structures were used as the basis for single point calculations of energy and of other properties at B3LYP/6-31G* level, using NBO at this level. Two sets of calculations were done:

1. *The cavity between the pore and the selectivity filter:* The starting protein coordinates were taken from the 3Lut pdb structure (38), which includes the hydrogens, added by normal mode analysis, and optimized with 50 molecules of water added; the coordinates of the water were optimized along with the protein. The upper limit on the system included the threonine of the TVGYG sequence, with water, and extended to the backbone of valine beyond. The K_v_1.2 cavity contains many more than the 12 water molecules of KcsA; we have here included some water that started, and sometimes remained after optimization, slightly outside the cavity, for the 50 molecule total. The calculation includes 870 atoms, 720 from the protein, plus the K^+^ ion. Six cases have been optimized: one without the ion, then one with the ion at a position approximately 2 Å above the PVPV gate where it optimizes (when the ion is placed at the level of the PVPV sequence, and not frozen, it moves, so that there is a barrier at this level—its energy is higher than the energy of the ion 2 Å further up). The ion is then frozen at the optimized position (labeled 0 Å), and at positions 2, 4, 6, and 8 Å above this, and then the rest of the system optimized. Of the protein atoms, 178 atoms near the periphery were frozen; it was >6 Å from the nearest frozen atom to a water oxygen, for all atoms; near the center of the cavity, >8 Å. Freezing peripheral atoms was required to make up for the absence of other atoms that would have held these atoms in place. All side chains that could come in contact with the ion or a water molecule were free. No water molecules were frozen; the potassium ion positions given are frozen with respect to the protein frozen atoms. Each calculation is separate, so if there were significant random errors, one would not expect that the energy values would produce a coherent result. One additional calculation with a second ion at the bottom of the cavity was also done, and gave a result slightly closer to the X-ray structure with the ion near the crystallographic ion position, and the stretch at the upper end that is described in results gone; in this case, the ion to protein atom distances everywhere were within approximately 0.5 Å of the X-ray value.
2. *The top of the cavity to the bottom half of the selectivity filter:* A set of calculations with 457 atoms were done that deleted the entire bottom of the cavity, and added the full valine above the threonine of the SF, plus the glycine above that. Three calculations each were done with two K^+^, and with two Na^+^, to compare hydration, and understand selectivity. The initial ion positions were 1) at approximately the 8 Å position of the ion in the 871 atom calculation, with the second ion near the second SF position—conventionally S3 (configuration A in Table 4 and Fig. 4); 2) with one ion near the conserved threonines (position S4) with the second ion several Å above that (near S3) (configuration B), 3) one ion near the 8 Å position of the cavity, the other between S3 and S4, with a water molecule in S4 (configuration C). In each case a water molecule separates the ions, and Fig. 4 (A→B→C) traces a path for K^+^ from the 8 Å position to the position between S4 andS3, together with the path for a water molecule that precedes it. The 457 atom calculation includes 13 water molecules. In this case, neither ions nor water was frozen; the different starting positions led to different minima, which did not coincide with the starting positions, but ions did not move far enough to find a different minimum. 52 protein atoms were frozen. Because the ions were not frozen, these ion positions correspond to an energy minimum. Methods and calculation level were the same as for the first calculations.

## RESULTS 1): ION IN THE CAVITY BETWEEN GATE AND SELECTIVITY FILTER

1) For the first set of calculations, Fig. 1 shows the system with the ion in each calculated position, except that the case with no ion is not shown. Fig. 1 shows a group of four water molecules just below the SF (the “basket” mentioned above) under the threonines of the SF TVGYG sequence, with hydrogen bonds to the carbonyl oxygens of the threonines; the hydrogens in the basket water molecules are only shown in Fig. 1A. In Fig. 1E, these water molecules begin to interact with the ion, and in Fig. 1F they interact directly with the ion.

**Fig. 1:**
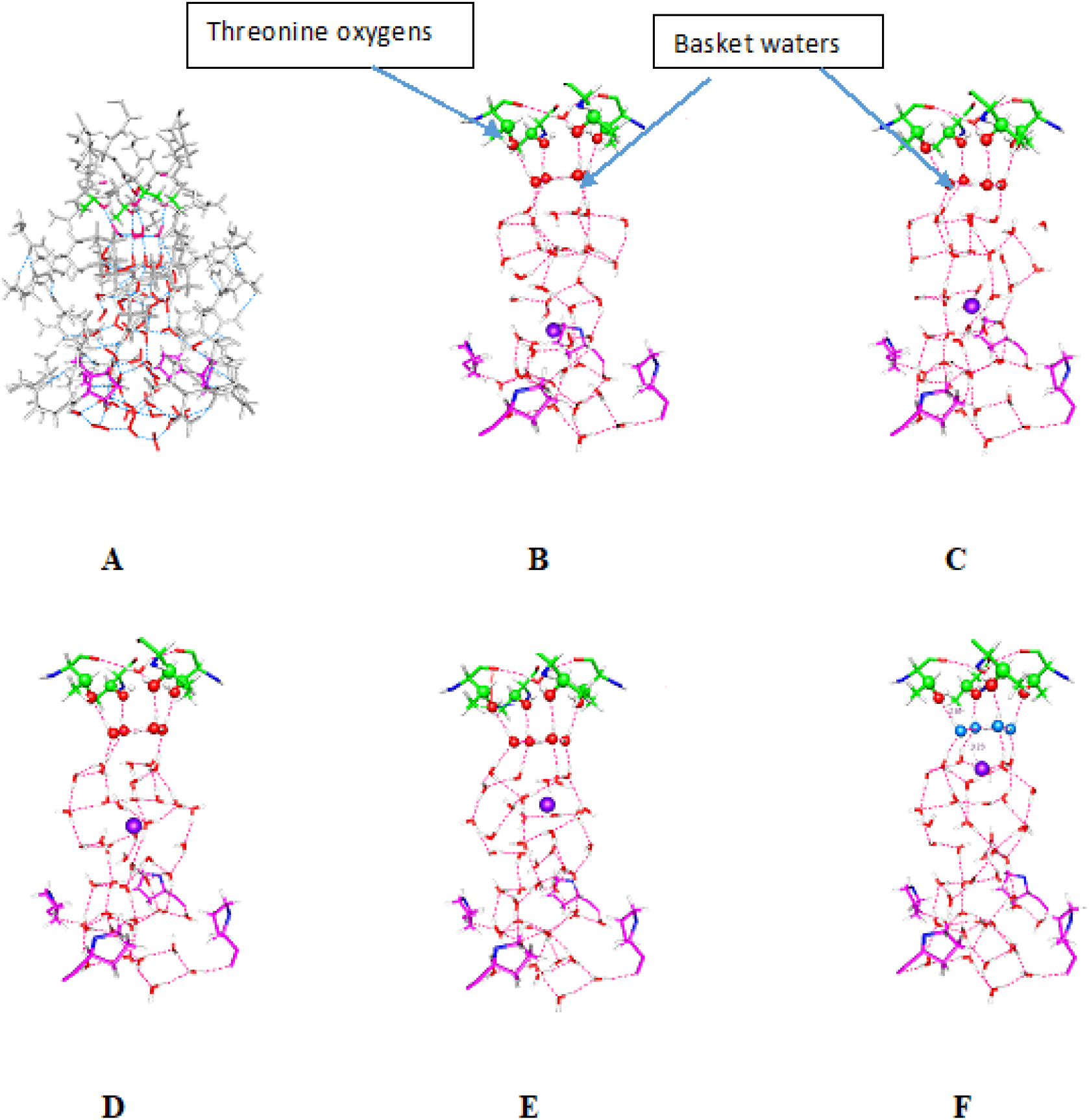
Water molecules in the cavity with the ion in five positions: Fig. 1A includes the protein, and is shown in a smaller scale than for figs 1B through 1F, for which all but the bottom of the SF (top, threonine carbons green), and (bottom) one proline (purple) of the PVPV sequence at the gate, are removed for clarity. The ion positions are at approximately 2 Å above the PVPV region (B), and then at 2 Å intervals above that (C through F). Dotted lines show hydrogen bonds, small red circles water oxygens (cavity), light blue, Fig 1F only, basket water interacting with the K^+^; large red circles, special oxygens, as labeled.

There is electron density transfer to the ion, so that the water becomes very slightly positive, as the ion loses charge. In addition, as we see below (Fig. 3), when the ion is at or below the center of the cavity, there is not a great deal of difference in the total dipole moment of the water molecules whether the ion is present or not, but above the center the difference is large. Finally, when the ion is above the position in Fig. 1F, it largely disrupts the basket, and is partially solvated with the oxygens from the threonines (see second calculation set).

The calculation shows an energy minimum with the ion in the center of the cavity, consistent with the fact that an ion is found approximately there in the crystal structure (Fig. 2). The minimum we calculate is at a position within approximately 0.5 Å of the crystallographic position, by comparison with three atoms at or below this point; the calculation is done at 2 Å intervals, limiting resolution. However, atoms near the top of the calculation are ≥2 Å higher with respect to the ion at the center than the X-ray distance, as though the cavity had expanded by perhaps 2 Å. Calculations in one case with a second ion below the central ion and near the PVPV gate level, however, give back the X-ray dimensions to within 0.5 Å. This suggests that the dimensions of the cavity, and the positions of the water, are affected by the presence of the ions in the cavity. Table 1 shows the distances for this case, where the optimized structure reproduces the X-ray distances.

**TABLE 1.** Comparison of calculated to X-ray distances of the cavity central ion, with second ion present

| Atom/amino acid* | O (OH)/Thr374 | CG1/Val399 | CG2/Val406 | N(ring)/Pro407 |
| --- | --- | --- | --- | --- |
| Distance (Å) calculated | 6.52 | 6.12 | 8.65 | 11.88 |
| Distance (Å) X-ray | 6.85 | 6.06 | 8.22 | 11.44 |

\*the ion distance is determined to these specific atoms on the designated amino acid

**Fig. 2.**
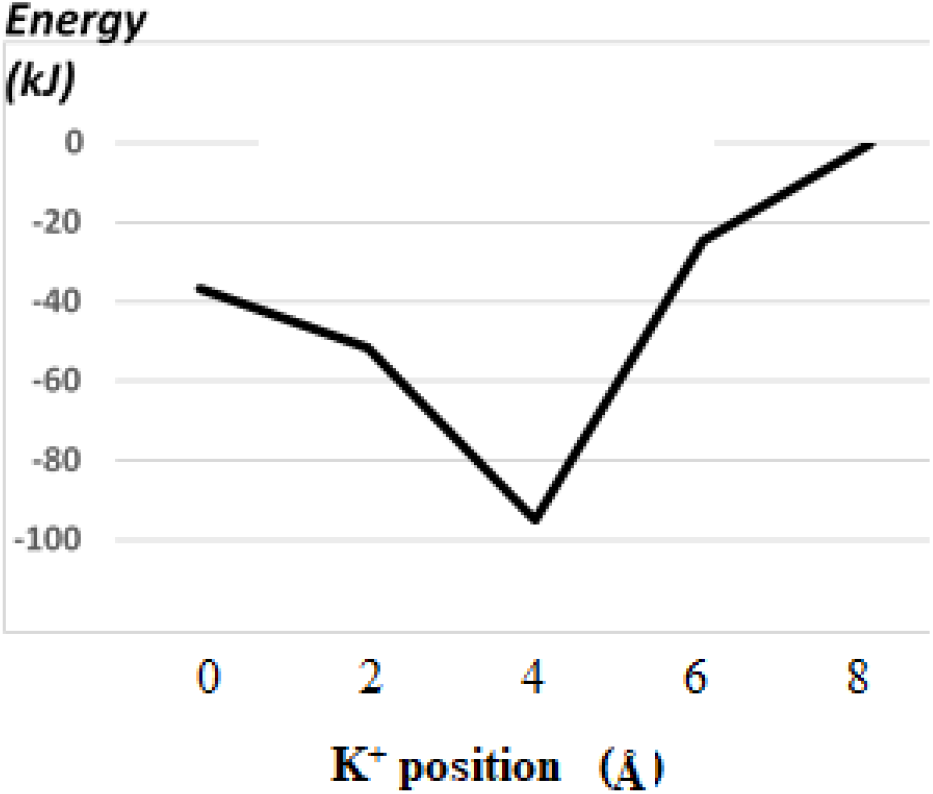
Energy of the entire system as a function of position of the K^+^ ion

The depth of the center energy minimum in Fig. 2, from the single ion calculations, seems appreciably too large. However, this single ion energy curve is sufficient to show that the ion is stable at the center of the cavity when a second ion has not yet entered the cavity. Including a second ion is almost certainly going to prove necessary to get complete agreement with experimental energy requirements for the ion to move forward in the pore; the one two ion calculation so far gives the coordinates with one ion at the center position, not the complete curve, so the well depth is not yet determined. The single ion results are necessary first, or the two ion results could not be interpreted, as the interaction effects could not be isolated. Furthermore, the single ion states may well exist as transient states. The one ion results are close to the X-ray structures, and the one calculation with a second ion, very close (Table 1). As the K^+^ position is not that close without the second ion, the agreement in Table 1 is significant; the calculation allows for disagreement.

We expect that the dipole moments of the water should follow the ion, rotating as the ion moves up. Figure 3 shows the vertical (*i.e.*, parallel to pore axis) component of the dipole moment of the water molecules only (no protein contribution), with and without the ion, for the five positions. The water molecule coordinates are identical for the two curves in the figure, showing the large effect of the ion’s charge in the upper region of the cavity on the charge distribution in the water molecules. Table 2 gives all components of the dipole moment. The other two components show the effect of the ion to be smaller, but not entirely negligible.

**Fig. 3:**
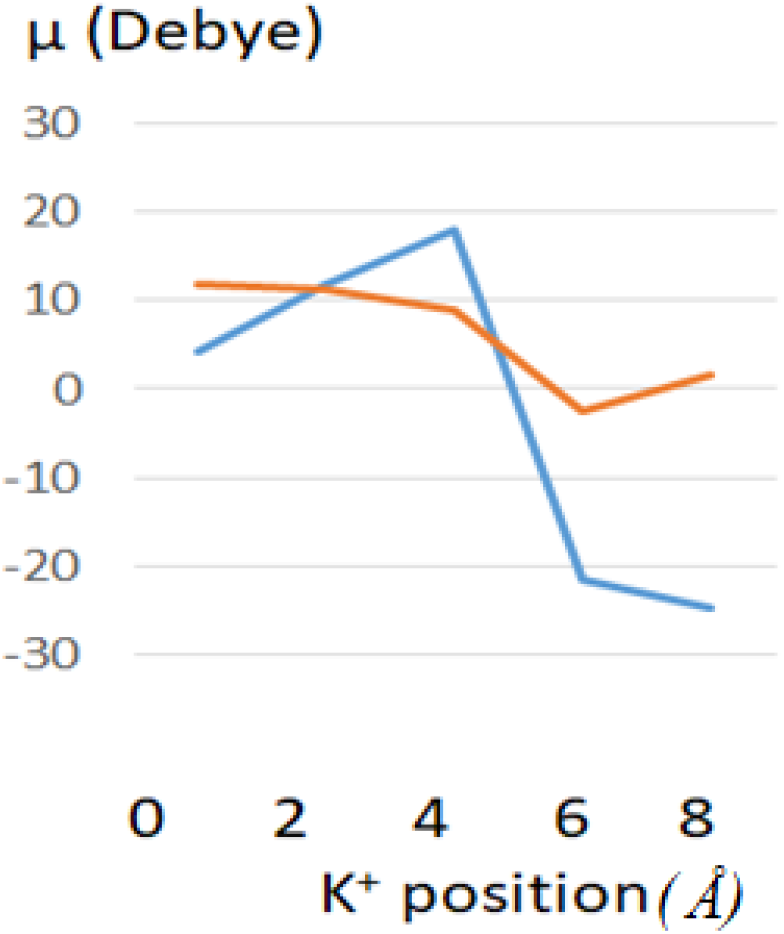
The “vertical” (along the pore axis) component of the dipole moment of the water molecules alone (blue) and the water molecules when the K^+^ ion is present (orange) for the five positions of the ion. The water molecules have the same coordinates in both lines; only the presence of the ion is different. The sign is positive when the positive pole of the dipole is up. No protein atoms are included.

**TABLE 2.** Dipole moment of the water in the cavity for five ion positions; x is the vertical component, others orthogonal

| Position (Å)<br>component | 0 | 2 | 4 | 6 | 8 |
| --- | --- | --- | --- | --- | --- |
| x (without $K^+$)* | 4.35 | 11.83 | 18.08 | -21.46 | -24.78 |
| y “ | 1.57 | 0.55 | 4.89 | 0.24 | 1.61 |
| z “ | -0.76 | 12.48 | 6.54 | 6.74 | 1.75 |
| total “ | 4.67 | 17.20 | 19.84 | 22.50 | 24.90 |
| x (with K<sup>+</sup>)* | 12.00 | 11.29 | 8.80 | -2.53 | 1.72 |
| y “ | -1.80 | -2.02 | 6.29 | 1.67 | 1.64 |
| z ” | -3.03 | 12.28 | 7.34 | 3.67 | 0.53 |
| total “ | 12.50 | 16.81 | 13.07 | 4.76 | 2.44 |

The interaction of the water with the protein, and, especially, the ion, leads to the rearrangement of the total dipole, especially the component parallel to the pore axis, together with the charge transfer to the ion of some additional electron density from the water, while some of the water molecules acquire a net charge. Much of the electron density transfer may in part be better thought of as from the protein, but transmitted through the water. The *difference* in the vertical component of the water molecules dipole moment with an ion in the upper part of the cavity, 18.93 D for the ion at 6 Å, 26.50 D for the ion at 8 Å, is large, and suggests that the water molecules, between polarization and charge transfer, are greatly different when the ion is present in these positions.

As water coordinates are identical, the dipole difference is a product of charge transfer and polarization, along with the charge on the ion itself. With the ion, the vertical component of the dipole is positive even when the ion is at the 0, 2, and 4 Å positions, so the majority of the dipole is due to the water molecules, not the ion; the ion should contribute in the opposite direction at positions 0 and 2. Although the ion shifts the center of charge, the fact that it makes much more difference when near the top suggests that it is not simply the charge on the ion that determines the dipole. The ion in the top two positions may make a large contribution to the overall charge distribution, in addition to that of the water molecules; in these positions the water alone has a large negative dipole, which is largely cancelled apparently by the positive charge on the ion.

Charge transfer to the ion lowers the charge on the ion, shown in Table 3. We note that the Na^+^ ion, which has a nominal 1s^2^2s^2^2p^6^ configuration, nevertheless accepts charge to an extent almost as great as does K^+^ with its 1s^2^2s^2^2p^6^3s^2^3p^6^ configuration; the NBO Natural Population Analysis (NPA) calculation assigns charges to the K^+^ 4s, 4p, and higher, orbitals, while Na^+^ would have to accept charge principally into 3s and 3p. The valence and Rydberg orbitals are not empty, as they would be if the charge were to be +1.00 on the ion. Charge transfer between water and ions has been found in several previous studies by a number of authors (39–41), and has been found to be appreciable. For a hydrated ion alone, the potassium ion charge has been reported as ≈0.87(40). The extent of charge transfer, however, depends on the links of the neighboring molecules from which the charge is drawn. Other studies show somewhat larger transfer, or show different methods of determining the extent of charge transfer (42–46). Huetz et al obtained results similar to ours in the SF, with charge transfer leaving charge a little less than 0.7 *e* on K^+^, slightly less than we found in some sections of the SF. In part this is because their results, and most of ours, do not include diffuse functions in the basis sets; the few cases we have done with the computationally expensive diffuse functions suggest that this would somewhat reduce the extent of charge transfer. The K^+^ charge was 0.775 without the diffuse functions (6-31G**), 0.802 with 6/31+G**, with the upper ion in Fig. 4A. This is not a large difference. However, when the ion was placed at the bottom of the structure (*i.e.*, at the top of the cavity, below the SF, near the 8 Å position in the 871 atom computation, but using the 457 atom SF computation), thus essentially in water, and some distance from protein, the K^+^ charge was 0.781 without diffuse functions, 0.872 with them. This is a substantial difference; the latter matches the result in the literature for water (40), while the former suggests that without the protein at least, it is necessary to have the diffuse functions. Charge transfer is substantial by any measure, and with the protein, there is a larger charge transfer. Surprisingly, the charge transfer to Na^+^ is not much different: With the diffuse functions, the ion charge in the upper position is 0.774 *e*, in the lower position, 0.891 *e*. Charge transfer changes as the ion moves. Water may be able to draw electron density from the protein, which in turn can be added to the K^+^. This suggests caution in quantitative interpretation of the results in Table 3. However, the principal qualitative conclusions, that there is substantial charge transfer, that the protein is important in this, and that it is position dependent, are unchanged. The effect is discussed at length by Weinhold and Landis, with an explanation of its origin in the back bonding of the bond orbitals (Chapter 2 of their book in particular)(47). MD simulations would not reflect this position dependence of charge transfer.

**Fig. 4:**
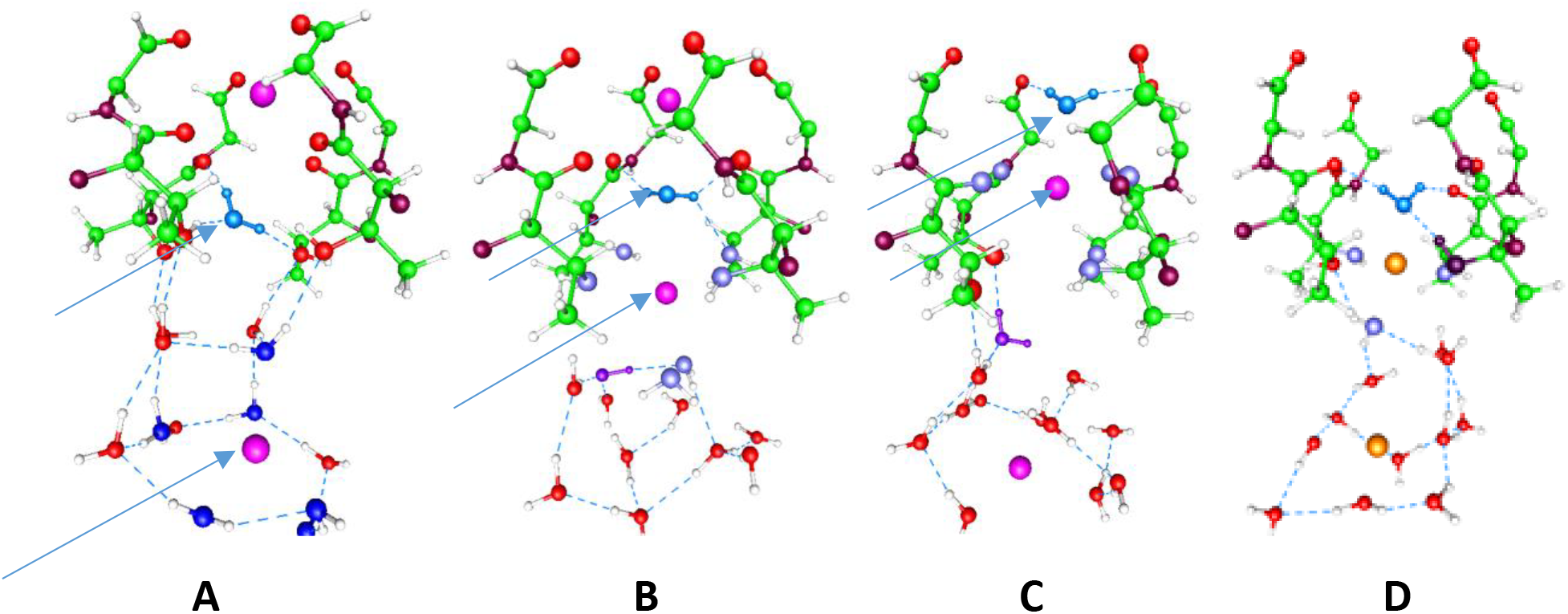
Subset of atoms cut from 457 atom calculations. Progress of a K^+^ ion and water molecule (arrows) from cavity (A) to the lowest position of the selectivity filter (S4, B) to the next position of the selectivity filter (S3, C). The one Na^+^ case is D. In A a second ion is in the S3 position at the top. In B that ion is still at S3, as the first ion moves up from the cavity (two steps may be needed). A water molecule (bright blue, arrow) separates the two ions. In C the ion is between S4 and S3, while the upper ion has moved up, out of the figure, and S3 is now occupied by the arrow water molecule (bright blue); at the bottom, an ion has moved into a position essentially the same as the lower arrow ion was in A, although with some rearrangement of water. In A, B, C, Carbon is green, nitrogen is dark red, hydrogen white, and oxygens which which do not solvate the arrow ion are red; oxygen atoms that are in the cavity and solvate the ion A are dark blue; protein oxygens that solvate the ion are pale blue (B and C). In B and C a key water molecule that rotates as the ion passes, reforming the S3 site, is purple; in C this is the water molecule that separates the two K^+^ ions. The upper Na^+^ ion in D (yellow) does not reach as high as the K^+^ in C, to which it should be compared. Two of the threonine hydroxyls are solvating the ion tightly, along with the water directly above it, only 2.27 Å away; that water is in turn strongly hydrogen bonded to the threonine hydroxyl shown in black on the right. One water below the ion also solvates it. The basket is broken. Differences between the lower Na^+^ in D and the lower K^+^ in C are not important.

**TABLE 3.** The charge and orbital occupancy on the K^+^ ion at the two extreme positions

| System(ion position, Å)* | 0(**water**) | 8(**water**) | 0(**total**) | 8(**total**) |
| --- | --- | --- | --- | --- |
| $K^+$ net charge | 0.706 | 0.674 | 0.741 | 0.672 |
| Fraction Valence electron density | 0.260 | 0.303 | 0.230 | 0.305 |
| Fraction Rydberg electron density | 0.027 | 0.023 | 0.030 | 0.026 |

\*0,8 = ion position, as defined in Fig. 1: **water**: $K^+$ orbital occupancy: 50 water molecules, no protein; **total**, $K^+$ orbital occupancy: entire system, with protein. NBO using B3LYP/6-31G\*\*

In Table 3, the K^+^ net charge is less than 1. The last two rows give the distribution of the electron density transferred to the ion; in essence, the second line shows the charge density in 3d + 4s + 4p orbitals, the last line to the sum over higher orbitals. If the net charge were a full +1.00, the last two rows would show 0.00 everywhere. The fact that this is not the case must be taken into account in understanding the behavior of the ions in the cavity, and the fact that the charge changes in different locations is something that must be accounted for in modeling ion transport in this system.

The NPA analysis showed some slight density in even 5s and 5p orbitals. When the protein is included in the calculation, there is 0.035 less charge transfer at the bottom of the cavity, but about the same at the top. The larger charge transfer difference as the ion moves up must affect the dipole of the system, The difference is not huge, but the energy involved is larger than k_B_T, hence sufficient to affect the transport of the ion. Much more important, as we discuss below with the calculation at the SF entrance, even the fairly small occupancy of the larger extension of the 5s and 5p orbitals may help K^+^ interact with all four threonines, while Na^+^ cannot. The effect of the protein on the charge transfer is significant, but by checking against the water-only background, we get a clearer picture of charge transfer.

## RESULTS 2): THE TRANSITION OF THE ION TO THE SELECTIVITY FILTER: THE STRUCTURE OF LOCAL ENERGY MINIMA

Finally, we need to know what happens as the ion penetrates the water basket immediately below, and hydrogen bonded to, the threonines, and then moves into the SF. Selectivity is one of the classic questions in ion channel function, with studies going back almost a half century(48). Calculations on a 457 atom subset of the pore (2 ions, 13 waters, 416 protein atoms), including the S4 (lowest) position of the SF, where the ion enters, plus most of S3, show the disruption of the basket, as well as the solvation of the ions by the oxygens from water and the oxygens of the threonines. We have found major differences in the way K^+^ and Na^+^ behave as they pass the water basket and bond to the threonine oxygens, with Na^+^ becoming trapped, largely by the hydroxyl threonines, while K^+^ passes these, interacts with carbonyl oxygens, and experiences essentially no barrier. This is shown in Table 4, while Fig. 4 shows the position of the ions, calculated from three starting positions. The K^+^ ion is apparently accompanied by the water to which it is effectively bound (above it, approximately in position S3 of the SF, in Fig. 4C), while migration of Na^+^ is trumped by an energetic wall; when the ion reaches the level of the threonine oxygens, there is a tightly bound water, at a distance of 2.7 Å ahead of it, while the ion is held by three hydroxyl oxygens from threonines. One Na^+^ calculation is shown (Fig. 4D) illustrating the point that the Na^+^ minimum is below that of the K^+^ position (Fig. 4C). The energies of the states through which the ions pass are also shown in Table 4; Na^+^ is faced with approximately a 14 k_B_T barrier going on from its position in Fig. 4D; experimentally, it takes approximately 300 mV to move Na^+^ through the channel(49). The only barrier to K^+^ is from Fig. 4 position (B) to position (C), and is only ≈5 k_B_T. A difference of 9kT corresponds to a factor of 8 × 10^3^, in the range of selectivity. However, the seemingly quantitative agreement is probably fortuitous. It shows that the calculation makes sense, but getting within one or two k_B_T is better than the expected accuracy of the calculation.

**Table 4.** Ion – oxygen distances*, bond orders and energy: upper K^+^ and Na^+^ in the last two positions

| Figure 4<br>correspondence<br>(4B,4C,4D) | Ion-oxygen<br>distances<br>to cavity<br>water | Ion-oxygen<br>distances<br>to<br>-OH<sup>1</sup> | Ion-oxygen<br>distances<br>to<br>-C=O<sup>1</sup> | Ion-oxygen<br>distance<br>to<br>water<br>above | Sum<br>of<br>bond<br>orders:<br>cavity<br>water | Sum<br>of<br>bond<br>orders<br>to<br>-OH | Sum<br>of<br>bond<br>orders<br>to<br>-C=O | Bond<br>order<br>to<br>water<br>above | $\Delta E^2$ (kJ<br>above<br>lowest<br>position) |
| --- | --- | --- | --- | --- | --- | --- | --- | --- | --- |
| $\text{K}^+(\text{B})$ | 2.88,2.93 | 2.68,2.79<br>2.82,2.87 | --- | 3.04 | 0.22 | 0.24 | --- | 0.26 | -33 |
| $\text{K}^+(\text{C})$ | --- | 3.22,3.29<br>3.95,4.70 | 2.89,2.96<br>2.85,3.08 | 2.75 | --- | 0.02 | 0.35 | -0.06 | -23 |
| $\text{Na}^+(-)\#$ | 2.30,2.40<br>2.44,3.75 | 2.40,2.65<br>3.93,4.02 | --- | 2.24 | 0.46 | 0.41 | --- | 0.11 | +35 |
| $\text{Na}^+(\text{D})$ | 2.40 | 2.29,2.47<br>2.88,2.90 | --- | 2.27 | 0.06 | 0.21 | --- | 0.02 | +1 |

Table 4 shows stark differences between Na^+^ and K^+^ in solvation, bonding, and energy. Na^+^ does not get to where it connects to the carbonyls at all. In the Na^+^(-) case we see the asymmetry of the bonding to the hydroxyls: 2 short distances, 2 quite long distances. K^+^ is not entirely symmetric itself, with one long distance. However, K^+^ goes higher, connecting fairly symmetrically to the carbonyls in (C), while Na^+^ cannot do this. Also, K^+^ goes down in energy to get to position (C), while Na^+^ has a large barrier. This is compensated to some extent in the next step, but, as Na^+^ never really reaches a comparable position, this is not a real comparison. Finally, we note that the water above the Na^+^ is quite close in the final position, 2.27 Å, but that water does not bond to the ion. Instead, it bonds to a water, shown in black in Fig. 4D, forming a lid over the ion. The ion in the previous position (Na^+^(-)) is strongly bonded to water below, and to two –OH groups. K^+^ is very different. In position (B) the bonding above and below is roughly equal, and in position (C) most of the bonding is to the carbonyls, and even that is less strong than that attaching the Na^+^. Taken together, Fig. 4 and Table 4 provide strong evidence for a series of steps that produces selectivity.

Also in these calculations; we again see the water molecules near the threonines at the bottom of the SF form a basket. A second ion may lower the barrier to moving the ion ahead of it to the SF, which would agree with the knock-on mechanism in a way, not by repelling the ion in the cavity center, but by rearranging the water. Direct ion-ion interaction would be likely to repel the incoming ion; the hydration states of the ions make the energy gradient favor the progress of the lead ion through the cavity while not repelling the incoming ion. For the K^+^ ions, not only does the barrier almost disappear, but the ion-ion distance drops to 5.57 Å. As shown in Fig. 4, the K^+^ and water form what appears to be a complex ion, which is fairly stable.

Fig.4 also shows the K^+^ ion has a coordination number of essentially 7, especially in part B. The number is not stable as the ion moves, and it has been characterized as dynamic by some authors(50, 51). Others have also considered the coordination number to be fundamental in selectivity(52)

A Na^+^ ion loses four-fold symmetry at the bottom of the SF. Table 4 shows the distances of the ions to the relevant oxygens of hydrating waters and carbonyl and hydroxyl for the three positions of each ion. The loss of symmetry with Na^+^ is evident from both Table and Figure. Also, Fig. 4 shows three positions of the K^+^ ions with its solvating oxygens, compared to the one position of the Na^+^ ion in which it is effectively trapped, which makes clear the difference between the two in the way they relate to the oxygens that solvate them; the asymmetry of the Na^+^ binding means that there are two strong interactions with oxygens of the threonine and two less strong. Dzidic and Kebarle showed that the hydrating waters for both Na^+^ and K^+^ as an isolated species bound water molecules sequentially, until 6 were reached, with monotonically decreasing energy (53), so the inequivalence of even nearly symmetric bonds may be expected. The Na^+^ coordination number is less than that of K^+^; in Fig. 4D it is essentially four, with two others, more distant, only possibly weakly relevant to coordination.

*Why is threonine so strongly conserved at the bottom of the SF*? The SF for this class of K^+^ channels starts with the sequence TVGYG over an evolutionary range encompassing viruses (2) and almost all prokaryotes and eukaryotes. The discussion above suggests a reason for the importance of this. There are three amino acids with side chains containing hydroxyls, serine, threonine, and tyrosine. Serine and threonine have different side chain lengths, and tyrosine’s properties are significantly different. The discussion above (Fig. 4 and Table 4) shows Na^+^ solvated by threonine hydroxyls, while K^+^ is solvated less tightly by the carbonyls. Table 4 shows the relevant distances from the ion. Threonine is unique in being able to do this; serine is sterically incapable of holding Na^+^, while tyrosine is completely different. A second threonine is very frequently paired with the one at the bottom of the SF (in 3Lut, T374 at the bottom of the SF, T373 paired with it in K_v_1.2, and a comparable pair is found in the majority of other channels of this type). T373 helps anchor the SF threonine, T374. The T374 hydroxyl hydrogen bonds to the T373 carbonyl oxygen (call these TT hydrogen bonds) under normal conditions, when K^+^ is present. When the K^+^ is below the four water basket, there are 3 TT hydrogen bonds, while with a K^+^ essentially at the level of T373, above the basket, there are 8 such bonds, which should be a more rigid structure, helping to insure correct solvation for K^+^.

So far there is no complete quantum calculation of the TVGYG sequence (the closest was Huetz et al (26), which gave a calculation that was QM/MM, and was not concerned with all aspects of the problem) but it is possible to hypothesize the main features of the part we have not calculated qualitatively; Na^+^ having already been selected out, the final GYG only has to let the K^+^ and accompanying water pass. The tyrosine –OH may be helpful in solvating the ion, while the glycines, lacking a side chain, allow space for the ion and water to pass. Calculations on this section are beginning, which should make the complete sequence of states clear.

## SUMMARY

1. The ordering of the water molecules around an ion passing through the cavity of the pore of the K_v_1.2 channel produces an energy minimum when the ion is at the center of the cavity, with the calculated position within 0.5 Å of the position of the ion in the crystal structure for the lower section, ≈ 2Å for the upper section. One calculation with two ions corrects this, with the ion position almost exactly that in the X-ray structure.
2. Charge transfer of electron density from the water to the ion is large, and there is an indirect contribution from the protein. The charge on the ion drops significantly below 1.00, to 0.8 or less in some positions.
3. Before the ion reaches the threonines at the bottom of the SF, four molecules of water form a “basket”, hydrogen bonded to the threonines; this is disrupted as the ion enters this position. In this position, earlier and present results show Na^+^ differs significantly from K^+^, losing symmetry, and being bound tightly enough to the protein to prevent it from moving with normal electric fields.
4. The sum of the water molecules net dipole is significantly differently oriented when an ion is included in the calculation, even when the water coordinates are identical, implying significant charge shift without structural change.
5. Two potassium ions in the lowest positions in the SF form what appears to be a complex ion that has a surprisingly low energy, and that allows the ions to progress to the next position with relatively little additional energy. The ion plus water pair moves from the lowest S4 position to S3 in a concerted fashion. This is not the case with two Na^+^ ions, which form configurations that lead to a major barrier to further progress into the SF.
6. It will be necessary to do more two ion calculations to gain a detailed quantitative understanding of the progress of the ion through the SF, especially of the interactions of the ions by way of the water, but the most important properties of the ion transport are at least qualitatively understood from the single ion results.

## AUTHOR CONTRIBUTIONS

AMK and MEG designed research, carried out research, and interpreted results. MEG wrote the paper, and AMK prepared the figures.

## ACKNOWLEDGEMENTS

This research used resources of the Center for Functional Nanomaterials, which is a U.S. DOE Office of Science Facility, and the Scientific Data and Computing Center, a component of the Computational Science Initiative, at Brookhaven National Laboratory under Contract No. DE-SC0012704. This research also used the High Performance Computation facility at City University of New York, funded by NSF.

